# Ancestry and Genetic Associations with Bronchopulmonary Dysplasia in Preterm Infants

**DOI:** 10.1101/263814

**Authors:** Dara G. Torgerson, Philip L. Ballard, Roberta L. Keller, Sam S. Oh, Scott Huntsman, Donglei Hu, Celeste Eng, Esteban G. Burchard, Roberta A. Ballard

## Abstract

Bronchopulmonary dysplasia in premature infants is a common and often severe lung disease with long term sequelae. A genetic component is suspected but not fully defined. We performed an ancestry and genome-wide association study to identify variants, genes and pathways associated with survival without bronchopulmonary dysplasia in 387 high-risk infants treated with inhaled nitric oxide in the Trial of Late Surfactant study. Global African genetic ancestry was associated with increased survival without bronchopulmonary dysplasia among infants of maternal self-reported Hispanic White race/ethnicity (OR=4.5, p=0.01). Admixture mapping found suggestive outcome associations with local African ancestry at 18q21 and 10q22 among infants of maternal self-reported African American race/ethnicity. For all infants, the top individual variant identified was within the intron of *NBL1*, which is expressed in mid-trimester lung and is an antagonist of bone morphogenetic proteins (rs372271081, OR=0.17, p=7.4 × 10^−7^). The protective allele of this variant was significantly associated with lower nitric oxide metabolites in the urine of non-Hispanic white infants (p= 0.006), supporting a role in the racial differential response to nitric oxide. Interrogating genes upregulated in bronchopulmonary dysplasia lungs indicated association with variants in *CCL18*, a cytokine associated with fibrosis and interstitial lung disease, and pathway analyses implicated variation in genes involved in immune/inflammatory processes in response to infection and mechanical ventilation. Our results suggest that genetic variation related to lung development, drug metabolism, and immune response contribute to individual and racial/ethnic differences in respiratory outcomes following inhaled nitric oxide treatment of high-risk premature infants.

## INTRODUCTION

Bronchopulmonary dysplasia (BPD) of premature infants is currently characterized by continuing requirement for supplemental oxygen and/or respiratory support at 36 weeks post menstrual age (PMA). BPD is the most common form of chronic lung disease in infants born prematurely and is associated with long-term respiratory morbidity, neurodevelopmental abnormalities, and death (35). The pathogenesis of BPD includes lung immaturity, with reduced pulmonary surfactant and low antioxidant and immune defenses, plus exposure to insults of hyperoxia, barotrauma from ventilator support, and infections that damage lung epithelium and elicit inflammation. Sequelae of this injury are arrested lung development, fibrosis and altered airway reactivity (7, 19, 24, 33, 35, 38, 47).

Therapeutic options for the prevention and treatment of BPD are limited and have not substantially affected the incidence of disease (reviewed in (26, 28)). For example, vitamin A treatment evokes a modest reduction of BPD but is not in general use, and caffeine reduces oxygen use and is routinely used for prevention of apnea. Postnatal dexamethasone therapy improves respiratory status acutely and decreases the incidence of BPD. However, longer courses of this therapy are associated with neurodevelopmental abnormalities. Inhaled nitric oxide (iNO) is used off-label in preterm infants to prevent BPD, however, the general efficacy of the drug has been brought into question (20).

The majority of studies evaluating the effectiveness of iNO have been performed in individuals with predominantly European ancestry (6). However, in the entire cohort of the Trial of Late Surfactant (TOLSURF) (60), and in a recent individual participant data meta-analysis across selected iNO trials (5), the incidence of BPD was significantly lower following treatment with iNO in infants of mothers who self-report as Black/African American ethnicity as compared to those who self-report as non-Hispanic White. Coupled to observed differences in levels of urinary NO metabolites in Black/African American vs. non-Hispanic White infants (8), these results suggest that response to iNO in terms of preventing BPD varies between racial/ethnic groups.

Although both the intrauterine and postnatal environment play an important role in BPD, twin studies have estimated the heritability between 50-80% (14, 39), suggesting a genetic contribution as well (29). Genetic studies of BPD have identified several candidate genes and pathways through genome-wide association studies (GWAS) (4, 29, 61) and exome sequencing (18, 40). However, none of the associations identified through GWAS have reached genome-wide significance, and replication of genetic associations has been problematic. This may in part be due to low statistical power given the relatively small sample size of each study (<1000 preterm infants), combined with the absence of a single genetic risk factor of large effect. Similarly, disease heterogeneity, including the potential for differences in the genetic architecture of BPD between racial/ethnic groups, and the specific definition of BPD used, may reduce statistical power. (4) However, pathway and gene-set enrichment analyses have identified candidates with high biological plausibility. (4, 40)

In this study, we performed a GWAS for survival without BPD in preterm infants in TOLSURF, which included infants of maternal self-reported African American, Hispanic, and non-Hispanic white race/ethnicity who all received iNO. We examine the effects of genetic variation at the level of individual variants, genes, and genetic pathways, and test the hypothesis that genetic ancestry at both the genomic and local scale is associated with survival without BPD in admixed populations.

## METHODS

### Study approval

Patient recruitment for the TOLSURF study was approved by the Institutional Review Boards at all participating sites including the University of California San Francisco.

### Study Subjects

TOLSURF was a masked, randomized, sham-controlled trial conducted in 25 US hospitals (ClinicalTrials.gov: NCT01022580). The study was designed to assess the effect of late doses of surfactant on BPD at 36 wk post menstrual age (PMA) in infants of 23-28 wk gestation who required intubation and mechanical ventilation between 7 and 14 days of age (9). A total of 511 infants were enrolled, and all received iNO (Ikaria, Hampton, New Jersey) according to the protocol followed in the NO CLD trial (10). BPD was assessed at 36 wk PMA by physiologic testing as described (10). There was no statistical difference in BPD incidence between control and surfactant-treated groups at 36 wk and the two groups were combined for this genetic study. Some infants were co-enrolled in the multi-center, observational Prematurity and Respiratory Outcomes Project (PROP) (52).

### Genotyping and Quality Control

DNA was extracted from tracheal aspirate cells from 454 infants whose parents consented for DNA collection using cells from up to five tracheal aspirate collections per patient. DNA was isolated using an AutoGeneprep 965 instrument (Autogen, Holliston, MA) by the manufacturer’s recommended standard protocol for human body fluids. In some cases, where protein contamination was evident, DNA was re-precipitated using 3 volumes of 100% Ethanol and 3M Ammonium acetate at a 3:1 ratio after incubation at −80°C overnight. Samples were initially quantified by Nanodrop (Thermofisher Scientific, Inc., Waltman MA) to access purity (A260/280) followed by analysis using the Agilent 2100 Bioanalyzer (Agilent, Santa Clara, CA) to more accurately access DNA quantity. The range of values for DNA concentration (ng/ul) ranged from 10-1750, median 130; total DNA/patient (ng) 130-4200, median 1600; A260/280 1.32-1.91, median 1.77; 429 samples were of suitable quality and quantity for genotyping.

Genotyping was performed on the Affymetrix Axiom LAT1 array (WorldArray 4) that contained >800,000 single nucleotide polymorphisms (SNPs) prior to quality control. SNPs were filtered based on call rates < 95%, and Hardy-Weinberg equilibrium p-values <10^−6^ using PLINK (53). Subjects were evaluated for call rates, consistency between genetic and reported sex, autosomal heterozygosity, and cryptic relatedness/genetic identity using IBD/IBS estimates in PLINK (53). In the case of multiples, one individual was selected at random to be included in the study.

### Statistical Analysis

Genomic levels of African ancestry were evaluated using ADMIXTURE (3) in a quasi supervised analysis assuming three ancestral continental populations of origin (K=3, African, European, and Native American). Windows were offset by a factor of 0.2, the cutoff for linkage was set to 0.1, and a constant recombination rate was set to 10^−8^ (bp)^−1^. The proportion of global African ancestry was compared between cases (BPD/death) vs. controls (survival without BPD) using logistic regression within infants with maternal self-reported African American ancestry and Hispanic ethnicity, adjusting for gestational age, sex, birth weight, and multiple gestation (yes/no). Local ancestry was inferred using LAMP-LD (11) in infants with maternal self-reported African American race/ethnicity using a two-population model. Unrelated CEU and YRI individuals from the HapMap were used as a reference to estimate global and local African ancestry.

Imputation of genetic variation from the phase 3 Thousand Genomes Project was performed using the Michigan Imputation Server (32), including ~ 79 million variants. Variants were then filtered for imputation quality scores > 0.3. Genetic association testing for survival without BPD was performed at both genotyped and imputed SNPs using logistic regression, adjusting for global genetic ancestry, gestational age, sex, birth weight, and multiple gestation. Analyses were performed within each racial/ethnic group using PLINK (53), then combined in a meta-analysis using METAL (64). Gene-based statistics were calculated using VEGAS (42) using genotyped SNPs, and intersected with a set of genes previously identified as being upregulated in BPD-dysregulated lungs (15). Pathway and gene-set analyses were performed using canonical pathways in IPA (Ingenuity Pathway Analysis (31)), and PANTHER (46) and MSigDB (56) using GREAT version 3.0.0 (45). Using GREAT, we assigned a foreground of gene coordinates with an association p>0.05 for survival without BPD, and a background of all gene coordinates for which a gene-based statistic was calculated (from VEGAS (42)).

Admixture mapping for local African ancestry was performed in infants with maternal self-reported African American race/ethnicity using logistic regression. Similar to association testing on individual variants, we performed association testing for the number of haplotypes of African ancestry at each genotyped SNP (homologous to association testing for the number of copies of the minor allele). Identical to our GWAS, we adjusted for global genetic ancestry, gestational age, sex, birth weight, and multiple gestation.

Measures of NO metabolites (NOx) including nitrate, nitrite, and nitrosylated compounds were made from the urine of 62 infants included in the current genetic study both before and following administration of iNO at 2-20 ppm as previously described (8). Briefly, urine was collected for 4-8 hours, and NOx were assayed according to (50) and normalized to creatinine to adjust for renal excretory function. NOx were measured at 3 different doses of iNO, including 2, 5, and 10-20 ppm. Genetic association testing at a single SNP was performed using linear regression to test for a correlation between genotype and values of NOx at a dose of 5 ppm. Values of NOx at 5ppm were selected for analysis because they are highly correlated to levels at 2 ppm, and more closely resemble a normal distribution as compared to 10-20 ppm.

For selected genes of interest, mRNA expression levels were obtained from a previous study that performed RNAseq on 3 specimens of human fetal lung of 23 wk gestational age (Gene Expression Omnibus, accession number GSE83888) (12).

## RESULTS

Following quality control, our study included a total of 795,465 genotyped SNPs and 387 unrelated infants; demographics by respiratory outcome is shown in Table 1 for 271 infants who died or had a diagnosis of BPD and 116 survivors without BPD. Overall, mean values for gestational age and birth weight were approximately 25 wk and 700 g, respectively, for this group of infants who still required intubation and ventilation between 7 and 14 days of age, representing a cohort at high risk for BPD. Within infants of maternal self-reported non-Hispanic white ethnicity (White), infants with BPD/death had a significantly higher respiratory severity score (RSS) upon study entry as compared to survivors without BPD, but had no significant difference in gestational age, birth weight, sex, and multiple gestations. Within infants of maternal self-reported Black/African American ethnicity (Black/AA), infants with BPD/death had significantly lower gestational age, lower birth weight, and higher RSS as compared to No BPD. These differences for infants with/without BPD are consistent with the known influence of immaturity and severity of early lung disease on BPD. No significant differences in clinical characteristics were observed between the two groups of maternal self-reported White Hispanic ethnicity (White Hispanic).

**Table 1.** Baseline characteristics of participants from the TOLSURF study included in the GWAS. P-values represent comparisons using a student’s t-test for continuous measurements (gestational age, birth weight, and RSS at entry), and a chi-square test for categorical (% male, % multiple gestation). Demographics are shown by maternal self-reported racial/ethnic group; data are mean plus standard deviation in parentheses. RSS=respiratory severity score.

|  | Non-Hispanic White: |  |  | African American: |  |  | Hispanic White: |  |  |
| --- | --- | --- | --- | --- | --- | --- | --- | --- | --- |
|  | BPD/<br>Death | No BPD | p-value | BPD/<br>Death | No BPD | p-value | BPD/<br>Death | No BPD | p-value |
| N | 136 | 41 | n/a | 82 | 51 | n/a | 28 | 14 | n/a |
| Gestational<br>Age (weeks) | 25.4<br>(1.3) | 25.2<br>(1.2) | 0.52 | 24.9<br>(1.0) | 25.4<br>(1.0) | 0.008 | 24.9<br>(1.3) | 25.5<br>(0.95) | 0.12 |
| Birth Weight<br>(g) | 712<br>(182) | 750<br>(165) | 0.21 | 640<br>(147) | 704<br>(145) | 0.015 | 703<br>(155) | 740<br>(210) | 0.57 |
| % Male | 59.6 | 51.2 | 0.44 | 56.1 | 45.1 | 0.29 | 57.1 | 35.7 | 0.33 |
| % Multiple<br>Gestation | 15.4 | 22.0 | 0.46 | 9.76 | 9.80 | 1.0 | 3.57 | 14.3 | 0.53 |
| RSS at Entry | 4.0<br>(2.1) | 3.1<br>(1.4) | 0.0008 | 4.0<br>(2.2) | 2.7<br>(0.94) | <0.0001 | 3.8<br>(2.8) | 3.3<br>(2.0) | 0.49 |

### Global Ancestry and Admixture Mapping

Individual proportions of genomic African ancestry were consistent with expectations given maternal self-reported race/ethnicity (Fig. 1, A and B). Specifically, Black/AA infants had a higher degree of African ancestry (median=85% [range=40-100%]) as compared to White Hispanic infants (median=6.3% [range=1.2-63%]).

**Fig. 1.**
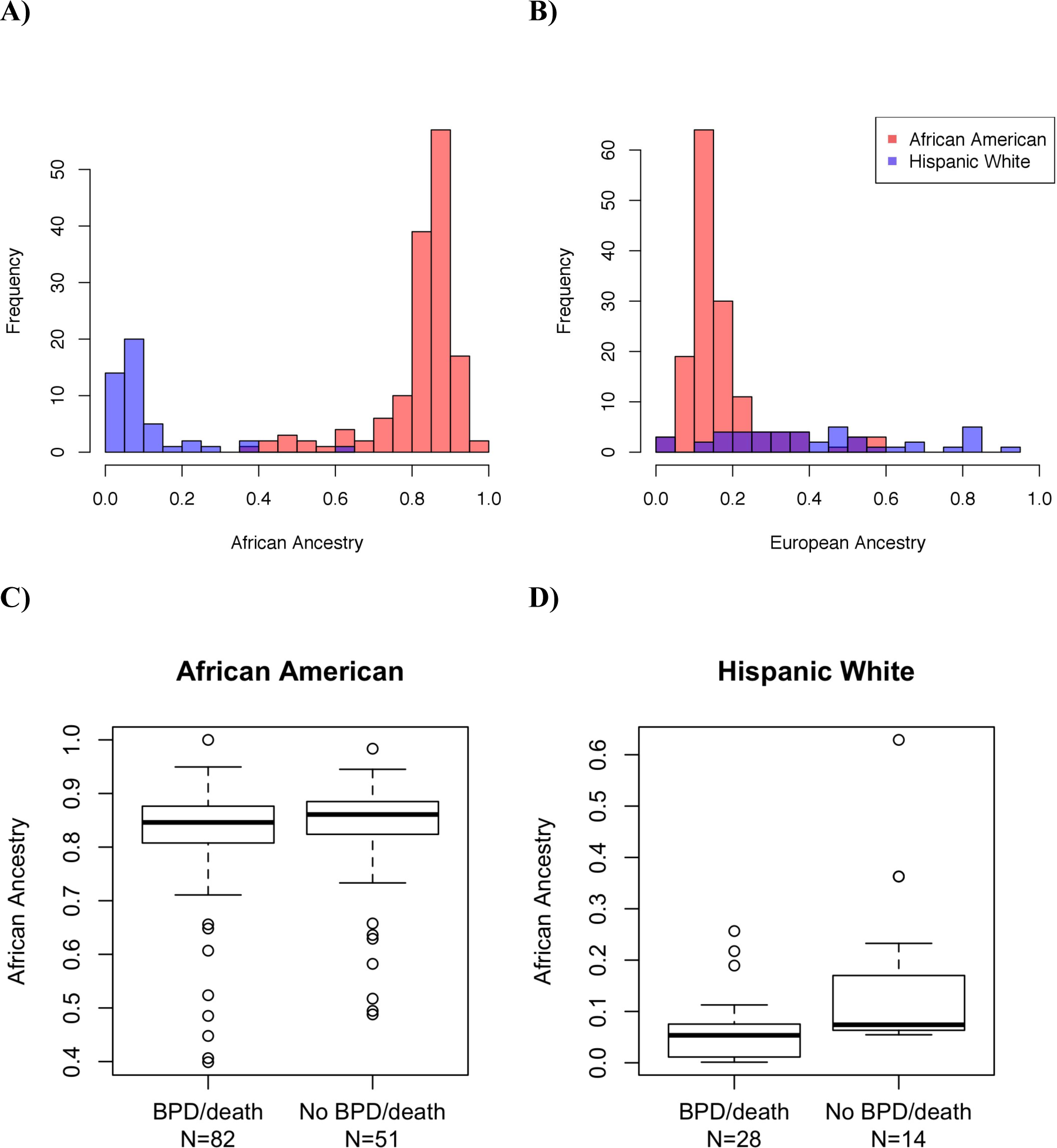
Global ancestry proportions and survival without BPD. The proportion of global African **(A)** and European **(B)** ancestry in preterm infants participating in the TOLSURF study by maternal self-reported race/ethnicity. Global ancestry was inferred using ADMIXTURE. Boxplots comparing global African ancestry and survival without BPD in infants of maternal self-reported Black/African American race/ethnicity **(C)** (logistic regression: p=0.97, β=−0.015 ± 0.37), and Hispanic White race/ethnicity (D) (p=0.01, β=−1.5 ± 0.60).

Global African ancestry was not significantly different between infants with BPD/death as compared to those surviving without BPD in Black/AA infants (beta=−0.015, se=0.37, p=0.97) (Fig. 1C). However, African ancestry was protective for BPD/death in White Hispanic infants (beta=−1.5, se=0.6, p=0.01) (Fig. 1D). Results were similar when all covariates were excluded. African ancestry was further compared at individual loci in Black/AA infants using logistic regression (i.e. local ancestry, or admixture mapping), and top associations were observed at 10q21 where African ancestry was protective for BPD/death (p=4.4x10^−4^, OR=0.17) and 18q21 where African ancestry was risky for BPD/death (p=2.7x10^−4^, OR=4.6) (Fig. 2). The estimated number of independent ancestry blocks was determined to be 478, and thus neither of the admixture mapping peaks was statistically significant following Bonferroni correction (alpha=1.0x10^−4^).

**Fig. 2.**
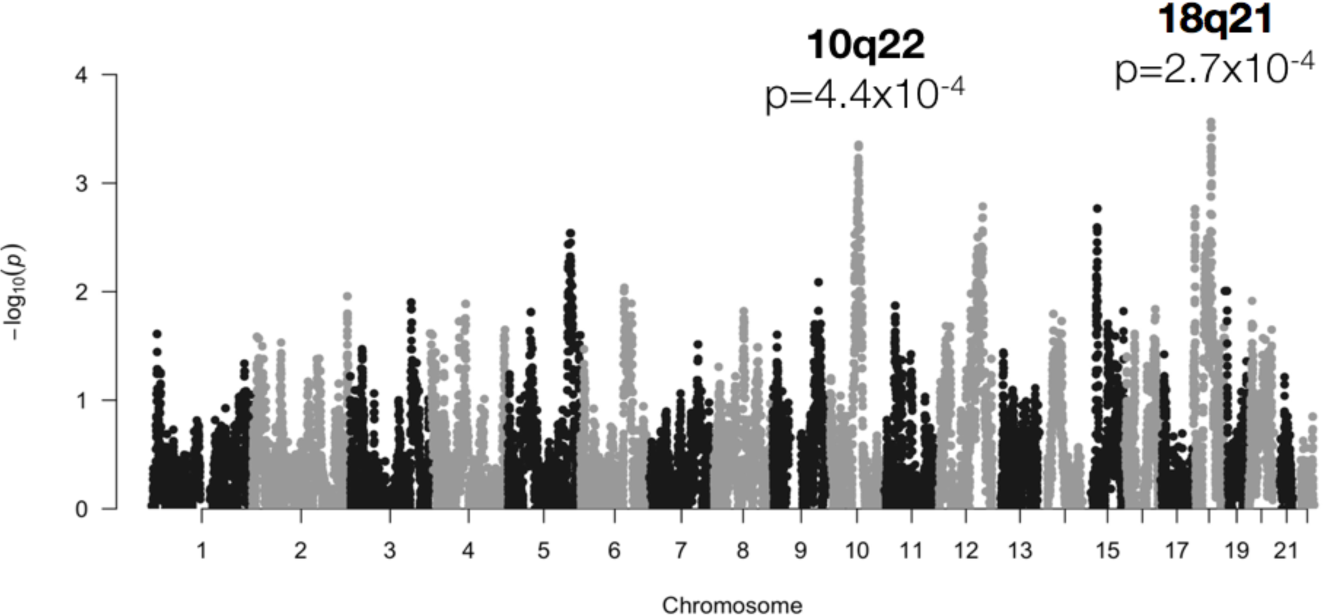
Results of admixture mapping comparing local African ancestry and survival without BPD in 133 infants with maternal self-reported Black/African American race/ethnicity (82 cases, 51 controls). Top associations were observed at 10q21 (OR=0.17, p=4.4x10^−4^) and 18q21 (OR=4.6, p=2.7x10^−4^).

### Genome-Wide Association Study (GWAS) and Gene-based Comparisons

Following genotype imputation, we tested the entire cohort for an association with survival without BPD at 8.8 million individual variants, adjusting for global genetic ancestry, gestational age, sex, birth weight, and multiple gestations. No individual variant was genome-wide significantly associated with BPD (all p-values > 5x10^−8^, Fig. 3). However, the top association was observed at a variant within the intron of NBL1 (rs372271081, p=7.4x10^−7^) (Fig. 4A). The minor allele was protective for BPD (OR=0.17), and showed a similar effect within each racial/ethnic group (Table 2). *NBL1* and two additional genes within the same region (*CAPZB* and *MINOS1*) were expressed in fetal lung at 23 wk gestation (Fig. 4B). Furthermore, the minor allele at rs372271081 was significantly associated with decreased urinary NOx in White infants, but was not significant in Black/AA or White Hispanic infants (Table 3, Fig. 5A). Notably, the protective allele for BPD at rs372271081 is at a somewhat higher frequency in populations with African ancestry (Fig. 5B).

**Fig. 3.**
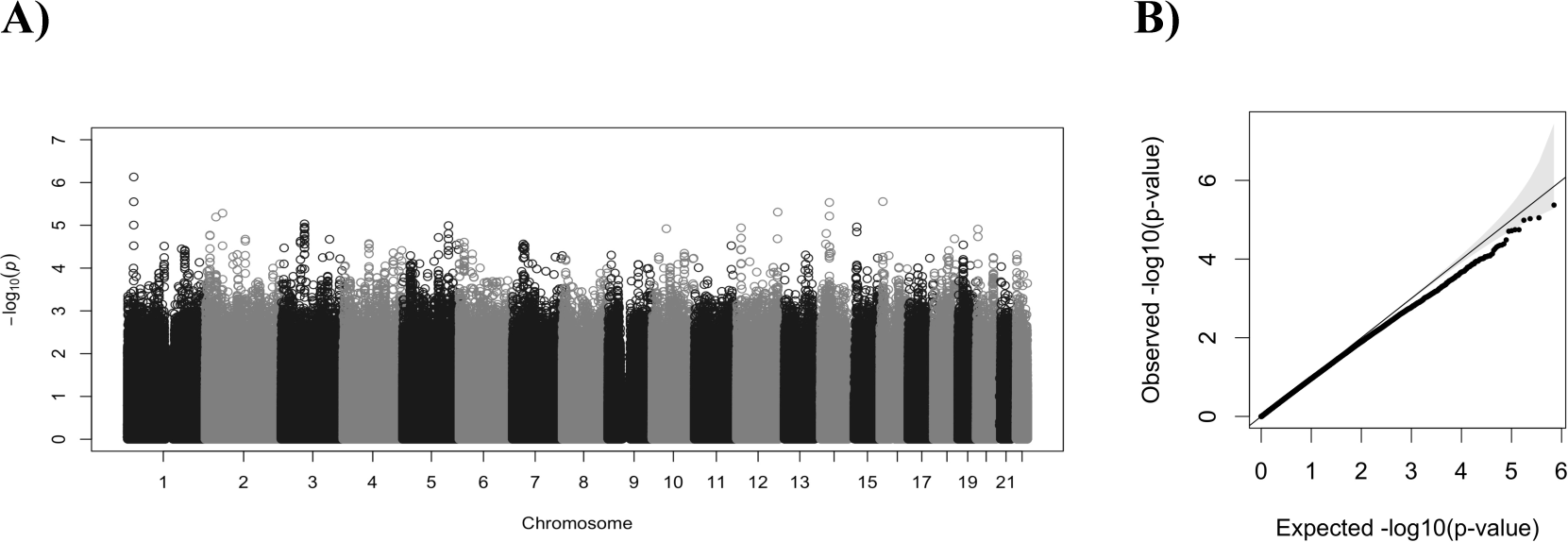
Manhattan plot (A) and quantile-quantile plot (qq plot, B) showing the results of a weighted meta-analysis for survival without BPD across three maternal self-reported racial/ethnic groups, including Non-Hispanic White (136 BPD/death infants, 41 no BPD), Black/African American (82 BPD/death infants, 51 no BPD), and Hispanic White (28 BPD/death infants, 14 no BPD).

**Fig. 4.**
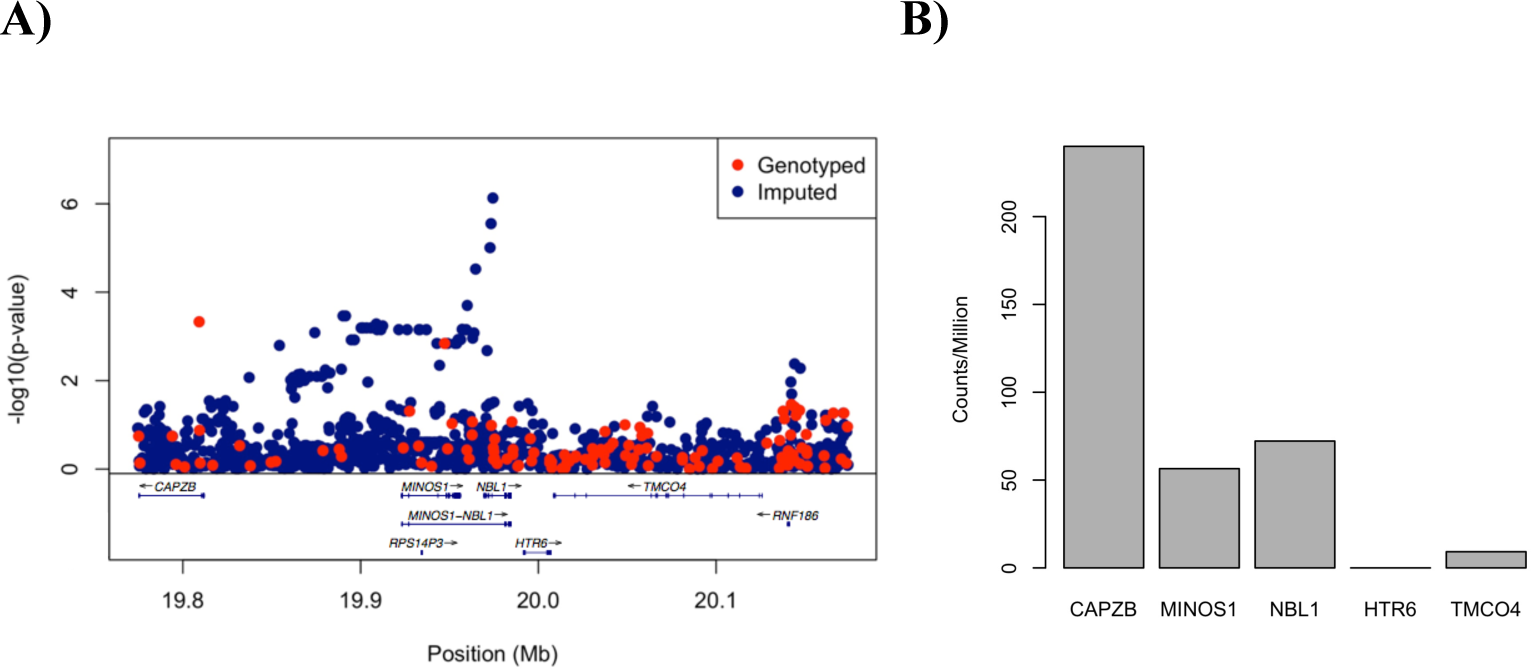
LocusZoom plot of the region flanking the top association at rs372271081, an intronic variant of *NBL1* (A). Expression of genes by RNAseq within this locus in fetal lung at 23 wk gestation (B).

**Fig. 5.**
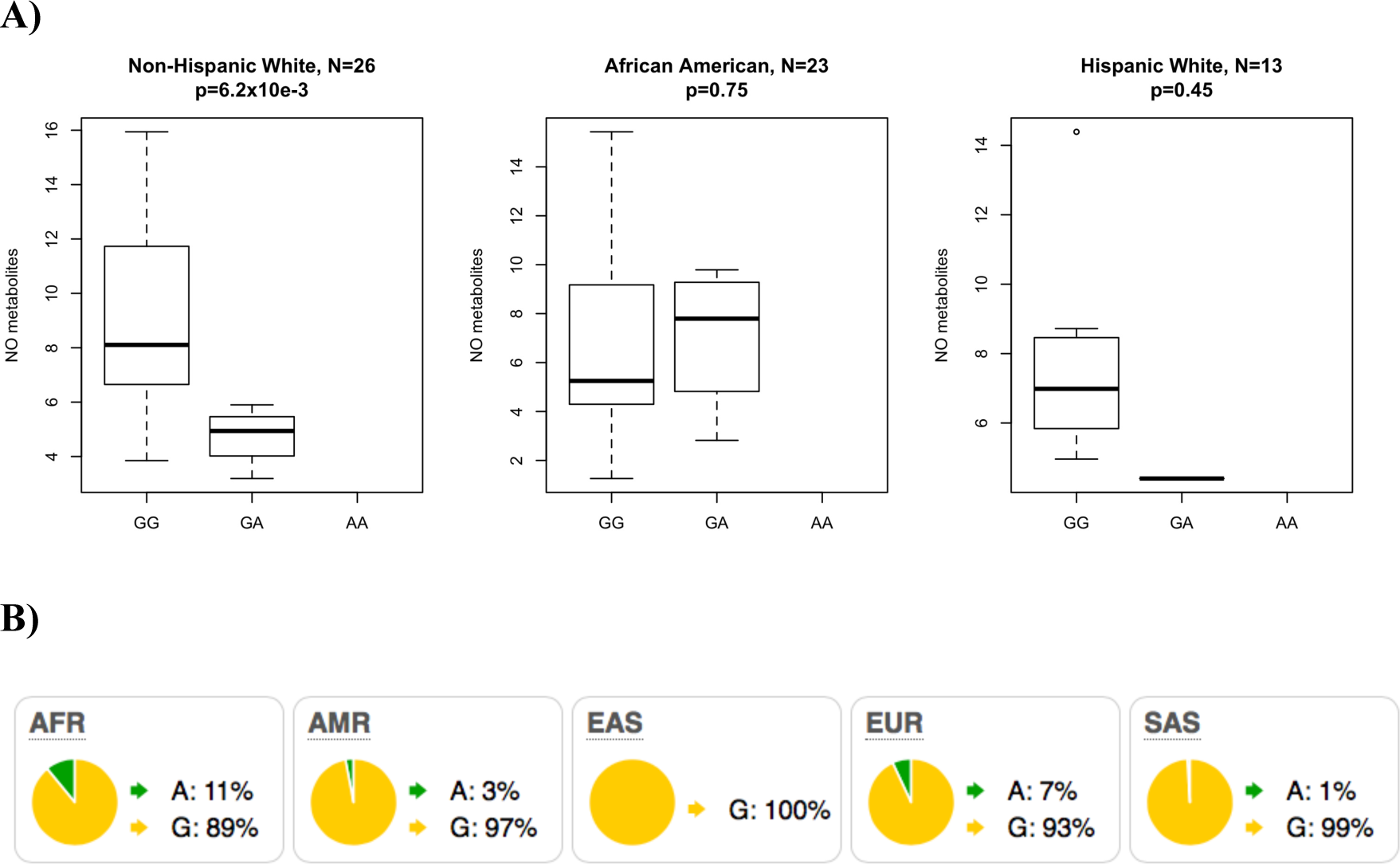
**(A)** Boxplot showing levels of urinary NO metabolites by genotype at rs372271081 in preterm infants following treatment with inhaled nitric oxide at 5ppm. **(B)** Frequency of rs372271081 in populations from the phase 3 1000 Genomes Project. AFR=African, AMR=Admixed American, EAS=East Asian, EUR=European, and SAS=South Asian.

**Table 2.** Results of tests of association at rs372271081 for survival without BPD using logistic regression, and urinary NO metabolites following iNO treatment using linear regression. Results are shown with respect to the minor allele (A), which trends as protective for BPD in three populations, and is significantly associated with lower urinary NO metabolites in infants of maternal non-Hispanic White race/ethnicity. Beta=regression coefficient/effect size, SE=standard error, CI=confidence interval.

|  | Survival without BPD: |  |  |  | Urinary NO metabolites: |  |  |  |
| --- | --- | --- | --- | --- | --- | --- | --- | --- |
| Population | Frequency in Cases (N) | Frequency in Controls (N) | Odds Ratio | P-value | N | Beta (SE) | 95% CI | P-value |
| Non-Hispanic White | 0.040 (136) | 0.12 (41) | 0.30 | $6.2 \times 10^{-3}$ | 26 | -5.3 (1.7) | (-8.7, -1.9) | $6.2 \times 10^{-3}$ |
| African American | 0.055 (82) | 0.19 (51) | 0.25 | $6.9 \times 10^{-4}$ | 23 | 0.74 (2.3) | (-3.8, 5.2) | 0.75 |
| Hispanic White | 0.018 (28) | 0.11 (14) | 0.15 | 0.070 | 13 | -1.9 (2.3) | (-6.5, 2.7) | 0.45 |

**Table 3.** Genetic variants associated with survival without BPD at p<10^−6^ in a meta-analysis across three racial/ethnic groups. For loci with multiple SNPs at p<10^−6^ only a single SNP with the smallest p-value is included in the table. OR=odds ratio, NHW=non-Hispanic White (136 BPD/death infants, 41 no BPD), AA=African American (82 BPD/death infants, 51 no BPD), HW = Hispanic White (28 BPD/death infants, 14 no BPD), Meta=meta-analysis (total 246 BPD/death infants, 106 no BPD).

| Chr | Position (hg19) | SNP | Allele | Annotation | NHW OR | AA OR | HW OR | Meta OR | Meta p-value |
| --- | --- | --- | --- | --- | --- | --- | --- | --- | --- |
| 1 | 19974397 | rs372271081 | A | intron, *NBL1* | 0.19 | 0.10 | 0.17 | 0.17 | $7.42 \times 10^{-7}$ |
| 2 | 14648908 | rs10193074 | G | intergenic | 0.26 | 0.25 | NA | 0.26 | $4.17 \times 10^{-6}$ |
| 2 | 33777089 | 2:33777089 | C | intron, *RASGRP3* | 0.39 | 0.17 | 0.22 | 0.28 | $6.41 \times 10^{-6}$ |
| 2 | 54980799 | 2:54980799 | G | intron, *EML6* | 0.33 | 0.78 | 0.44 | 0.40 | $5.20 \times 10^{-6}$ |
| 2 | 105035900 | rs4851694 | T | intergenic | 3.8 | 8.7 | NA | 4.3 | $5.92 \times 10^{-6}$ |
| 2 | 105039687 | rs2889323 | C | intergenic | 3.8 | 8.7 | NA | 4.3 | $5.92 \times 10^{-6}$ |
| 2 | 105091271 | rs6543256 | G | intron, *LOC150568* | 3.2 | 9.0 | NA | 4.1 | $7.24 \times 10^{-6}$ |
| 3 | 74073182 | rs1949931 | G | intergenic | 0.38 | 0.082 | 0.44 | 0.39 | $9.25 \times 10^{-6}$ |
| 10 | 134044152 | rs60417571 | T | intron, *STK32C* | 0.19 | NA | 0.27 | 0.21 | $3.06 \times 10^{-6}$ |
| 12 | 131048872 | 12:131048872 | CTG | intron, *RIMBP2* | 0.44 | 0.11 | 0.41 | 0.39 | $4.91 \times 10^{-6}$ |
| 14 | 47459909 | rs8016110 | A | intron, *MDGA2* | 2.9 | 43 | 2.5 | 3.5 | $2.92 \times 10^{-6}$ |
| 16 | 8834085 | rs75055007 | A | intron, *ABAT* | 0.30 | 0.061 | 0.28 | 0.26 | $2.79 \times 10^{-6}$ |

To increase statistical power, we combined the results of association testing of individual variants within known genes to create a single gene-based statistic. No individual gene was significantly associated with BPD following Bonferroni correction for 17,670 tests (the number of genes tested, alpha=2.8x10^−6^) (Table 4). However, by restricting our comparisons to 21 candidate genes whose expression is dysregulated in BPD lungs, variation in *CCL18* was significantly associated with BPD (p=0.0011). This gene is expressed at a low level in 23-wk human fetal lung (0.31±0.10 cpm). None of the genes implicated from Li et al. (40) were significantly associated with BPD (minimum p=0.0018, *ADCY8*), nor were 11 NO-related candidate genes (Table 5) with reported associations to human disease (minimum p=0.18, *KALRN*).

**Table 4.** Top genes associated with survival without BPD in a meta-analysis across racial/ethnic groups including 246 cases and 106 controls. Gene-based statistics were calculated using VEGAS, none of the genes were statistically significant following Bonferroni correction for the total number of genes examined (N=17,671).

| Chr | Gene | Number of SNPs | p-value |
| --- | --- | --- | --- |
| 5 | *RICTOR* | 17 | $4.5 \times 10^{-5}$ |
| 4 | *MED28* | 6 | $1.8 \times 10^{-4}$ |
| 12 | *IL23A* | 8 | $2.2 \times 10^{-4}$ |
| 19 | *ZNF492* | 14 | $2.5 \times 10^{-4}$ |

**Table 5.**
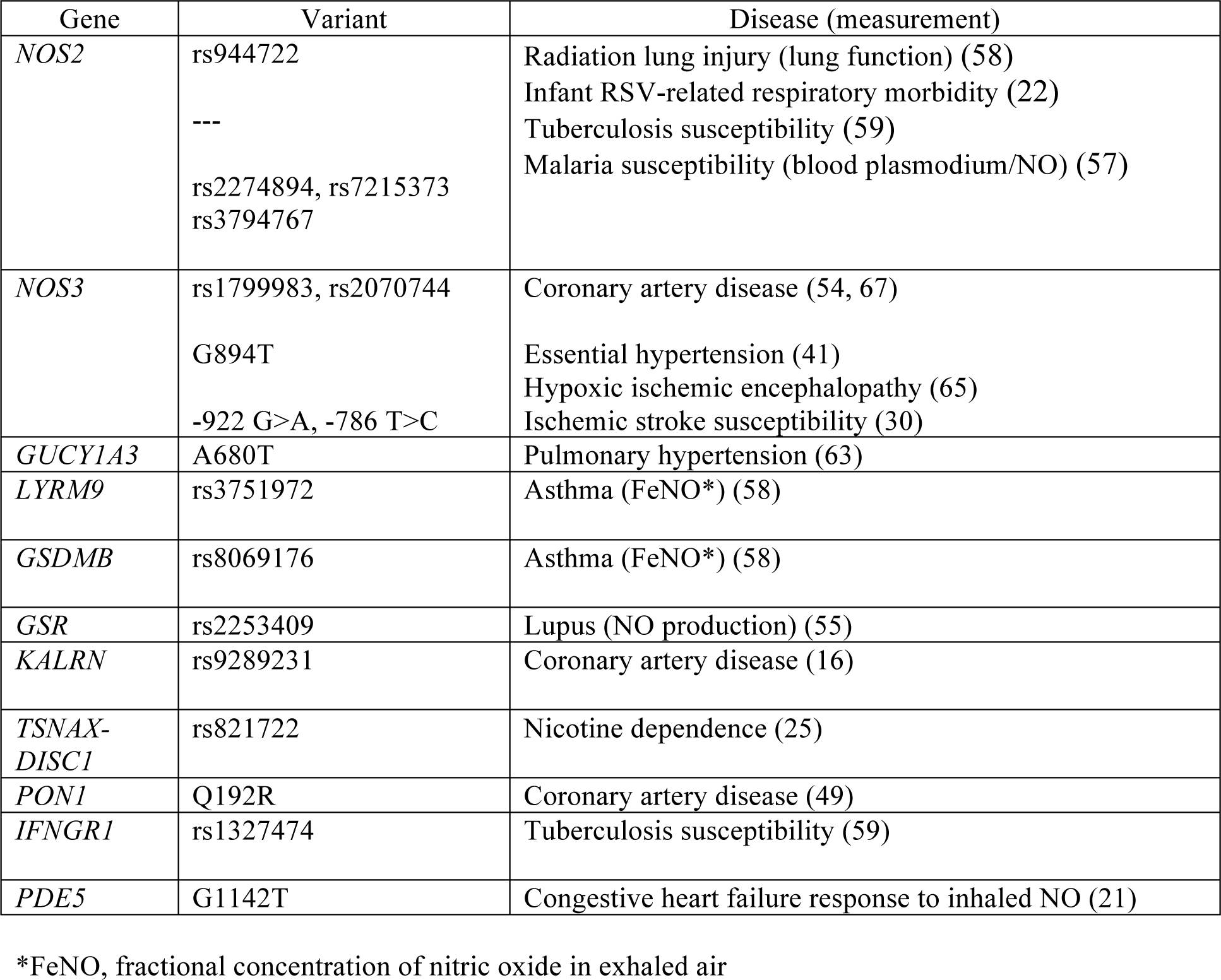
List of NO-related candidate genes/variants previously associated with disease.

### Pathway Analysis

Pathway analysis can be a powerful means to identify an enrichment of genes with marginal signals of association on their own, but which function in a similar biological pathway. Pathway analysis was performed using GREAT (45) on 1,024 genes with gene-based p-values < 0.05 as compared to 17,640 genes as a background. A total of five pathways/gene sets were identified with a false discovery rate (FDR) < 0.05 from the Panther and MSigDB databases; this group contains two pathways related to cancers, one related to immune function, one related to methylation marks, and one implicated in experimental lung injury (Table 6). The pathway of highest statistical significance (p=5x10^−12^) was ‘Genes within amplicon 1q21 identified in a copy number alterations study of 191 breast tumor samples”. Eight of the 11 genes in this pathway are expressed in human fetal lung at 23 wk GA, and none are regulated by glucocorticoids, which enhance fetal lung maturity. Biological functions of potential relevance to lung development, injury, and repair for these genes include tyrosine kinase receptor signaling pathway (EFNA4, RUSC1, SHC1, ADAM15), developmental processes (EFNA4, RUSC1, ZBTB7B, PBXIP1, SHC1, ADAM15, PYGO2) including angiogenesis (SHC1), NF-kappaB signaling (RUSC1, ZBTB7B), and sex steroid receptor signaling (PBXIP1, SHC1).

**Table 6.** Panther and MSigDB pathways showing a significant enrichment of genes associated with survival without BPD at p<0.05. A foreground set of 1,024 genes with association p-value < 0.05 was compared to a background set of 17,640 genes using GREAT. Gene-based association p-values were calculated using VEGAS. FDR=false discovery rate.

| Pathway/Gene Set | P-value | FDR | Fold Enrichment | Number of Genes |
| --- | --- | --- | --- | --- |
| **Panther:** |  |  |  |  |
| Toll receptor signaling pathway | $2.3 \times 10^{-4}$ | 0.035 | 3.1 | 13 |
| **MSigDb:** |  |  |  |  |
| Genes within amplicon 1q21 identified in a copy number alterations study of 191 breast tumor samples | $1.5 \times 10^{-15}$ | $5.1 \times 10^{-12}$ | 8.6 | 21 |
| Genes with low-CpG-density promoters bearing H3K4me3 marks in embryonic fibroblasts | $4.1 \times 10^{-7}$ | $6.8 \times 10^{-4}$ | 2.5 | 35 |
| Nearest neighbors of TAL1, based on the close agreement of their expression profiles with that of TAL1 in pediatric T cell acute lymphoblastic leukemia | $6.7 \times 10^{-6}$ | 0.0076 | 5.0 | 11 |
| Genes up-regulated in lung tissue upon LPS aspiration with mechanical ventilation | $2.4 \times 10^{-5}$ | 0.020 | 2.2 | 32 |

Pathway analysis using Ingenuity Pathway Analysis (IPA) of 181 genes with gene-based p-values < 0.01 of 209 canonical pathways identified two with a significant enrichment of genes following a Bonferroni correction: agranulocyte adhesion and diapedesis (p=3.06x10^−5^) and granulocyte adhesion and diapedesis (p=1.22x10^−4^); genes in these two pathways are identical except for *MYL9* (Table 7). With the exception of *CLDN17*, all genes identified in these pathways are expressed in human fetal lung (12).

**Table 7.** Canonical pathways from Ingenuity Pathway Analysis (IPA) with a significant enrichment of genes with association p<0.01 for survival without BPD. Statistical significance was determined using a Bonferroni adjustment for 209 canonical pathways tested (alpha=2.39x10^−4^).

| Canonical Pathway | # of Genes (%) | Genes in Pathway with $p < 0.01$ | P-value |
| --- | --- | --- | --- |
| Agranulocyte adhesion and diapedesis | 9/181 (4.8%) | CCL3, CCL4, CCL17, CCL18, CCL22, CLDN17, CX3CL1, MYL9, RDX | $3.06 \times 10^{-5}$ |
| Granulocyte adhesion and diapedesis | 8/181 (4.4%) | CCL3, CCL4, CCL17, CCL18, CCL22, CLDN17, CX3CL1, RDX | $1.22 \times 10^{-4}$ |

## DISCUSSION

Unique aspects of our study are the patient population and rigorous assignment of BPD. All infants in TOLSURF were <28 wk gestation and were intubated at 7-14 days, representing infants with severe early respiratory failure and high risk for BPD as reflected by the occurrence of BPD/death in 68.5% of the total population (9). In addition, infants were enrolled from 25 different U.S. sites, providing both racial/ethnic and geographic diversity. The diagnosis of BPD was assigned on a physiologic basis using an oxygen/flow reduction challenge to establish a requirement for respiratory support. Thus, it is possible that some of our findings may be restricted to extremely premature infants with severe early respiratory disease. Other unique characteristics of our cohort are exposure to late surfactant treatment in approximately half of the infants, and the use of iNO for 3 wk in all infants. Although surfactant therapy transiently improved respiratory status it did not affect outcome at 36 wk PMA. iNO therapy likely influenced outcome in African American infants, but not Caucasians, and thus we examined NO metabolism as it relates to genetic associations with BPD in our study.

Higher genomic levels of African ancestry were associated with better respiratory outcome in iNO-treated infants with maternal self-reported Hispanic White race/ethnicity but not for infants with maternal self-reported Black/African American race/ethnicity. While the protective effect of African genomic ancestry in White Hispanic infants requires independent replication, our results suggest that a protective effect of African ancestry may be saturated at lower levels of ancestry than is present in the majority of Black/African American infants in the study, or may reflect the presence of differential gene-environment interactions. Specifically, in White Hispanic infants, the proportion of African ancestry ranged from 1.2-63% (median of 6.3%) as compared to 40-100% (median of 85%) in Black/African American infants.

Genetic ancestry does not form a direct causal relationship, but rather indicates differences in the underlying patterns of genetic variation in infants with/without BPD that differ by continental origin. If only small proportions of African ancestry are required for a protective effect, this would suggest a highly polygenic contribution to BPD distributed throughout the genome. Admixture mapping in Black/AA infants identified two suggestive, but not statistically significant peaks, at 10q22 and 18q21, whereby African ancestry was associated with both a decreased and increased risk of death/BPD, respectively. Therefore, admixture mapping further supports the hypothesis that the effect of ancestry is not limited to a single locus of large effect. It is possible the relationship between ancestry and BPD is restricted to infants receiving iNO, given that the incidence of BPD in many studies doesn’t vary between racial/ethnic groups in untreated infants (13); furthermore, prior studies have identified racial differences in endogenous NO levels or metabolism in infants (8) and adults (34, 36, 37, 44).

In our agnostic scan including ~9 million genotyped and imputed variants, no individual variant was genome-wide significantly associated with survival without BPD. This was not unexpected given our small sample size, and is consistent with prior GWAS that similarly failed to identify individual variants with large effects (4, 29, 61). Modest sample size is a limitation common to genetic studies of preterm infants, and thus there is a need to integrate additional biological measurements. Along this line, our top BPD-associated variant, rs372271081, was significantly associated with differences in NOx in White infants, whereby the protective allele for BPD was associated with decreased levels of NOx in the urine following treatment of iNO. However, we found no significant association between genotype and NO metabolites in Black/AA or Hispanic White infants, which may reflect the limited statistical power given the smaller sample sizes, the lower frequency of the allele, varying patterns of linkage disequilibrium, and/or the presence of genetic and environmental interactions.

Rs372271081 lies within an intron of Neuroblastoma 1, DAN Family BMP Antagonist (*NBL1*), which is a highly plausible candidate gene for contributing to BPD susceptibility via differential response to iNO. Numerous studies in mice indicate that the BMP pathway is important for lung development, including branching morphogenesis in early gestation and distal lung epithelial cell differentiation, alveolization and vasculogenesis in late gestation (17, 23, 62). The TGF-β/BMP signaling pathway is disrupted by hyperoxia (1), which is known to play a role in the development of BPD (1, 2). In humans, disrupted BMP signaling has been implicated in the pathogenesis of heritable pulmonary arterial hypertension and hereditary hemorrhagic telangiectasis (27, 48). Lastly, in addition to ligand inhibition of BMP, DAN family members are known to modulate wnt and VEGF signaling pathways that have a role in lung development and injury/repair (48). Overall, there is strong biological plausibility for a role of genetic variation in *NBL1* and respiratory outcome in iNO-treated infants based on 1) the critical role of BMP signaling in lung development and disease, 2) the mediation of BMP action via NO, 3) the expression of *NBL1* and BMPs in human fetal lung (35), and 4) the racial differences in BPD and NO metabolism (12).

Although *NBL1* has not been specifically implicated in prior GWAS/exome sequencing studies, genes involved in lung development are strong candidates for a role in BPD, which only occurs in immature lungs. For example, a common variant in *SPOCK2*, an extracellular matrix protein, was implicated in BPD through GWAS and found to be upregulated during lung alveolar development and after exposure to hyperoxia in rats (29). Furthermore, pathway analyses have implicated other genes involved in pulmonary structure and functions (40). Replication and both laboratory and functional validation are necessary to confirm a causal relationship of variants in *NBL1* and BPD in infants treated with iNO. Currently there are no other cohorts of premature infants treated with iNO with DNA samples available for validation studies.

We further performed hypothesis-driven tests of association with BPD using a set of 21 genes that are dysregulated in BPD lungs (15), and 11 genes in the nitric oxide pathway that are reported to have variants associated with disease (Table 5). First, we hypothesized that genes showing differential expression in BPD-dysregulated vs. control lungs may contain variants that contribute to survival without BPD. We found a significant association with genetic variation in a single gene - *CCL18*, a cytokine involved in the immune response that promotes collagen production in lung fibroblasts (43) and is associated with pulmonary fibrosis and interstitial lung diseases in adults (51, 66). Inflammation is known to be important in the pathogenesis of BPD, and anti-inflammatory therapy (dexamethasone) suppresses a variety of inflammatory mediators, including CCL18, and reduces BPD (12, 26). Second, because all infants in the study received iNO, we hypothesized that variation in genes in the NO pathway may contribute to differential response to iNO treatment as indicated by survival without BPD. Yet, no individual variant or candidate gene (based on known association with human disease) was significantly associated with survival without BPD following correction for multiple tests.

However, because differences in rates of BPD between racial/ethnic groups may be exclusively observed in infants treated with iNO (5, 60), we hypothesized that genetic variants that contribute to BPD may act through differential response to iNO. In support of this, the protective allele for BPD at rs372271081 is significantly associated with decreased NOx and is more common in populations with African ancestry. Several studies indicate reduced bioavailability of NO in African Americans vs Caucasians, likely in part due to increased oxidation of NO. In laboratory studies, release of NO from umbilical venous endothelial cells was substantially lower in African American vs Caucasian infants (36, 44). Levels of urinary NOx are lower in African American and Hispanic premature infants vs Caucasian infants regardless of iNO treatment, reflecting baseline differences in NO metabolism and thus bioavailability (8). In adults, African Americans are known to have increased frequency of hypertension and cardiovascular disease, and a NO-targeted medication (isosorbide dinitrates and hydralazine) is indicated therapy for heart failure specifically in African Americans (i.e., a racially directed therapy) (34, 37). However, further studies are needed to evaluate the contribution of rs372271081 to racial/ethnic differences in NO bioavailability and differential response to iNO.

Pathway analyses identified pathways and sets of genes that were significantly enriched for genes with association p-values < 0.05. Across IPA, Panther, and MSigDB datasets, a common theme that emerged was genes involved in immune function, including granulocyte and agranulocyte adhesion and diapedesis from IPA canonical pathways, toll receptor signaling pathway from Panther, and genes upregulated in response to LPS exposure and mechanical ventilation from MSigDB. These results suggest that variation in immune response, including recruitment of leukocytes and lymphocytes, contributes to survival without BPD.

Overall, our results for this cohort of iNO-treated, high-risk infants suggest that genomic African ancestry is protective for BPD, and that an intronic variant in *NBL1* may contribute to BPD via differential activity of the TGF-β/BMP pathway and production/metabolism of NO. Furthermore, we implicated variation in genes involved in the immune response, including *CCL18*, as contributing to differences in respiratory outcomes of preterm infants.

## AUTHOR CONTRIBUTIONS

D.G.T., P.L.B., R.L.K. E.G.B. and R.A.B. conceived and designed research; P.L.B., R.L.K., C.E., E.G.B., and R.A.B. performed experiments; D.G.T., P.L.B., R.L.K., S.S.O., S.H., D.H., C.E., and R.A.B. analyzed data; D.G.T, P.L.B., R.L.K., and R.A.B. interpreted results of experiments; D.G.T. prepared figures; D.G.T. and P.L.B. drafted manuscript; D.G.T., P.L.B., R.L.K., E.G.B. and R.A.B. edited and revised manuscript; D.G.T., P.L.B., R.L.K., S.S.O., S.H., D.H., C.E., and R.A.B. approved final version of manuscript.

## ACKNOWLEDGMENTS

We thank the TOLSURF Investigators, study coordinators, physicians, nurses, respiratory therapists and the families who participated in the TOLSURF study.

## AUTHOR NOTES

Address for reprint requests and other correspondence: Dara G. Torgerson, University of California San Francisco, MC 2530, 1700 4^th^ Street, San Francisco CA 94143.

## APPENDIX TOLSURF Investigators

Robin H. Steinhorn, Department of Pediatrics, Children’s National Medical Center, Washington DC

Catherine M. Bendel, Department of Pediatrics, University of Minnesota, Minneapolis MN

Ellen M. Bendel-Stenzel, Department of Pediatrics, Children’s Hospital and Clinics of Minnesota, St. Paul and Minneapolis MN

Sherry E. Courtney, Department of Pediatrics, University of Arkansas for Medical Sciences, Little Rock AR

Ramasubbareddy Dhanireddy, Department of Pediatrics, University of Tennessee Health Science Center, Memphis TN

Frances R. Koch, Department of Pediatrics, Medical University of South Carolina, Charleston SC

Mark L. Hudak, Department of Pediatrics, University of Florida College of Medicine, Jacksonville FL

Dennis E. Mayock, Department of Pediatrics, University of Washington, Seattle WA

Rajan Wadhawan, Department of Pediatrics, Florida Hospital for Children, Orlando FL

Nicolas F. Porta, Department of Pediatrics, Northwestern University Feinberg School of Medicine, Chicago IL

Victor J. McKay, Department of Pediatrics, All Children’s Hospital, St. Petersburg FL

Jeffrey D Merrill, Children’s Hospital Oakland Research Institute, Oakland CA

Eric C. Eichenwald; Department of Pediatrics, Children’s Hospital of Philadelphia, Philadelphia PA

William E. Truog, Department of Pediatrics, Children’s Mercy Hospital, Kansas City MO

Mark C. Mammel, Children’s Hospital and Clinics of Minnesota, St. Paul MN

Elizabeth E. Rogers, Department of Pediatrics, University of California San Francisco, San Francisco CA

Rita M. Ryan, Department of Pediatrics, Medical University of South Carolina, Charleston SC

David J. Durand, Children’s Hospital Oakland Research Institute, Oakland CA

T. Michael O’Shea, Department of Pediatrics, Wake Forest School of Medicine Winston-Salem NC

Dennis M. Black, Department of Epidemiology and Biostatistics, University of California San Francisco, San Francisco CA

In addition to the authors and other TOLSURF investigators, the following members of the TOLSURF Study Group participated in this study:

<u>University of California, San Francisco, San Francisco, CA:</u>

Suzanne Hamilton Strong, RN, Jill Immamura-Ching, RN, Margaret Orfanos-Villalobos, RN, Cassandra Williams, RN; Lisa Palermo MA

<u>Alta Bates Summit Medical Center, Berkeley, CA, and UCSF Benioff Children’s Hospital Oakland, Oakland, CA:</u> Dolia Horton, RRT, Loretta Pacello, RCP, April Willard, RN;

<u>Children’s Mercy Hospital, Kansas City, MO:</u>

Cheryl Gauldin, RN, Anne Holmes, RN, Patrice Johnson, RRT, Kerrie Meinert, RRT;

<u>Women and Children’s Hospital of Buffalo, Buffalo, NY:</u> Anne Marie Reynolds, MD, Janine Lucie, NNP, Patrick Conway, Michael Sacilowski, Michael Leadersdorff, RRT, Pam Orbank, RRT, Karen Wynn, NNP;

<u>Anne and Robert H. Lurie Children’s Hospital/Northwestern University, Chicago, IL:</u> Maria deUngria, MD, Janine Yasmin Khan, MD, Karin Hamann, RN, Molly Schau, RN, Brad Hopkins, RRT, James Jenson, RRT;

<u>Texas Children’s Hospital, Houston, TX:</u> Carmen Garcia, RN

<u>Stony Brook University Hospital, Stony Brook, NY:</u> Aruna Parekh, MD, Jila Shariff, MD, Rose McGovern, RN, Jeff Adelman, RRT, Adrienne Combs, RN, Mary Tjersland, RRT;

<u>University of Washington, Seattle, WA:</u> Elizabeth Howland, Susan Walker, RN, Jim Longoria, RRT, Holly Meo, RRT;

<u>University of Texas Health Science Center, Houston, TX:</u> Amir Khan, MD, Georgia McDavid, RN, Katrina Burson, RN, BSN, Richard Hinojosa, BSRT, RRT, Christopher Johnson, MBA, RRT, Karen Martin, RN, BSN, Sarah Martin, RN, BSN, Shawna Rogers, RN, BSN, Sharon Wright, MT;

<u>University of Florida College of Medicine, Jacksonville, UF Health Shands Hospital, and Wolfson Children’s Hospital, Jacksonville, FL:</u> Kimberly Barnette, RRT, Amanda Kellum, RRT, Michelle Burcke, RN, Christie Hayes, RRT, Stephanie Chadwick, RN, Danielle Howard, RN, Carla Kennedy, RRT, Renee Prince, RN;

<u>Wake Forest School of Medicine and Forsyth Medical Center, Winston Salem, NC:</u> Beatrice Stefanescu, MD, Kelly Warden, RN, Patty Brown, RN, Jennifer Griffin, RRT, Laura Conley, RRT;

<u>University of Minnesota Amplatz Children’s Hospital, Minneapolis, MN:</u> Michael Georgieff, MD, Bridget Davern, Marla Mills, RN, Sharon Ritter, RRT;

<u>Medical University of South Carolina, Charleston, SC:</u> Carol Wagner, MD, Deanna Fanning, RN, Jimmy Roberson, RRT;

<u>Children’s Hospitals and Clinics of Minnesota, St. Paul, MN:</u> Andrea Lampland, MD, Pat Meyers, RRT, Angela Brey, RRT;

<u>Children’s Hospitals and Clinics of Minnesota, Minneapolis, MN:</u> Neil Mulrooney MD, Cathy Worwa, RRT, Pam Dixon, RN, ANM, Gerald Ebert, RRT-NPS, Cathy Hejl, RRT, Molly Maxwell, RT, Kristin McCullough, RN;

<u>University of Tennessee Health Science Center, Memphis, TN:</u> Mohammed T. El Abiad, MD, Ajay Talati, MD, Sheila Dempsey, RN, Kathy Gammage, RRT, MBA, Gayle Gower, RN, Kathy James, RRT, Pam LeNoue, RN;

<u>All Children’s Hospital, St. Petersburg, FL:</u> Suzi Bell, DNP, Dawn Bruton, RN, BSN, CCRP, Michelle Beaulieu, DNP, Richard Williams, RRT;

<u>Florida Hospital for Children, Orlando, FL:</u> Robin Barron-Nelson, RN, Shane Taylor, RRT;

<u>Arkansas Children’s Hospital and University of Arkansas Medical Sciences, Little Rock, AK:</u> Carol Sikes, RN, Gary Lowe, RRT, Betty Proffitt, RRT.

<u>Clinical Coordinating Center:</u>

University of California San Francisco, Department of Pediatrics: Cheryl Chapin, Hart Horneman, Karin Hamann, RN, Susan Kelley, RRT, Karin Knowles, Nancy Newton, RN, MS.

<u>Data Coordinating Center:</u>

University of California San Francisco, Department of Epidemiology and Biostatistics: Eric Vittinghoff, PhD, Jean Hietpas, Laurie Denton, Lucy Wu.

<u>Data Safety Monitoring Board:</u>

Cincinnati Children’s Hospital Medical Center, Cincinnati, OH: Allan Jobe,MD, (Chair 2009-2010); UH Rainbow Babies and Children’s Hospital, Cleveland, OH: Avroy Fanaroff, MD (Chair 2010-2016); EMMES Corporation, Rockville, MD: Clemons; Boston University School of Public Health, Boston, MA: Leonard Glantz, JD; Wake Forest School of Medicine, Winston-Salem, NC: David Reboussin, PhD; Stanford University, Stanford, CA: Krisa Van Meurs MD (2009-2010); Johns Hopkins University, Baltimore, MD: Marilee Allen, MD (2010-2016); Women and Infants Hospital, Providence, RI: Betty Vohr, MD.

<u>Clinical Steering Committee:</u>

Department of Pediatrics, UCSF Benioff Children’s Hospital San Francisco, San Francisco CA: Roberta Ballard, MD, Philip Ballard, MD, PhD, Roberta Keller, MD, Elizabeth Rogers, MD, Nancy Newton, MS, RN; University of California, San Francisco, Department of Epidemiology and Biostatistics: Dennis Black, PhD; National Heart, Lung and Blood Institute (NIH/NHLBI): Carol Blaisdell, MD; UCSF Benioff Children’s Hospital Oakland, Oakland, CA: David Durand, MD, Jeffrey Merrill, MD, Jeanette Asselin, MS, RRT; University of Texas Health Science Center, Houston, TX: Eric Eichenwald, MD; Children’s Hospital and Clinics of Minnesota, St. Paul, MN: Mark Mammel, MD; Medical University of South Carolina, Charleston, SC: Rita Ryan, MD; Children’s Mercy Hospital, Kansas City, MO: WilliamTruog, MD.

